# HLA-DR MODULATION AND PD-1/PD-L2 CHECKPOINT SIGNALLING DEFINE A MECHANISTIC POTENCY AXIS FOR MESENCHYMAL STROMAL CELL IMMUNOSUPPRESSION

**DOI:** 10.64898/2026.05.01.722253

**Authors:** M Nikougoftar Zarif, Katia Lefsihane, N Khanlarkhani, L Sörvik, JF Talts, K Le Blanc, Kadri Nadir

## Abstract

Mesenchymal stromal cells exhibit potent immunomodulatory properties and are under active investigation for the treatment of immune-mediated disorders. However, their clinical translation is hindered by the lack of standardized potency assays. Here, we established a reproducible mixed lymphocyte reaction platform by systematically optimizing peripheral blood mononuclear cell donor composition, culture conditions, and co-culture ratios to define a robust activation window. Using this system, we compared bone marrow and adipose derived Mesenchymal stromal cells across independent donor batches.

Both sources effectively suppressed T cell proliferation, with the adipocyte derived source consistently showing greater inhibitory activity, while a conserved lower threshold of suppression was observed across both sources. Mesenchymal stromal cells reduced early (CD25^+^) and late (CD25^+^HLA-DR^+^) T cell activation, with downregulation of these markers emerging as a sensitive correlate of functional potency. Notably, bone marrow derived mesenchymal stromal cells exerted stronger suppression on late-stage activation and preferentially suppressed CD8^+^ T cell expansion.

Mechanistically, this immunosuppression was associated with modulation of the PD-1 pathway, characterized by decreased soluble PD-1, increased PD-L1, and induction of mesenchymal stromal cells derived PD-L2. PD-L2 levels inversely correlated with T cell proliferation, identifying a PD-1/PD-L2 regulatory axis linked to the cell’s potency.

These findings define a standardized and mechanistically informed potency assay framework for assessing mesenchymal stromal cell immunomodulatory function.

## Introduction

Mesenchymal stromal cells (MSCs) are multipotent progenitor cells present in multiple tissues that contribute to tissue repair and immune regulation. They are defined by the International Society for Cellular Therapy (ISCT) through plastic adherence, tri-lineage differentiation capacity and characteristic surface marker expression ^1^. MSCs exhibit potent immunomodulatory properties through the secretion of soluble factors and extracellular vesicles, making them promising candidates for treating immune-mediated diseases ^2,3^.

MSCs regulate immune responses by suppressing T cell activation and proliferation, modulating antigen presenting cells and promoting regulatory T cell induction ^2,4^. Both CD4^+^ and CD8^+^ T cell subsets are affected ^5^, with activation marked by early CD25 expression and later HLA-DR upregulation ^6,7^. Modulation of these activation states may provide informative functional readouts beyond proliferation-based assays.

Despite clinical success in indications such as graft versus host disease (GVHD) ^8^, a major challenge in the clinical development of MSCs based therapies is the lack of standardized potency assays. Regulatory agencies emphasize the need for robust, quantitative assays that reflect the mechanism of action relevant to the intended clinical indication ^9^. The mixed lymphocyte reaction (MLR) is widely used to measure MSCs-mediated immunosuppression^10^, but is highly sensitive to experimental variability, underscoring the need for optimized and reproducible assay platforms suitable for quality control ^11^.

Additionally, MSCs from different tissue sources exhibit functional heterogeneity, including differences in immunosuppressive capacity, molecular profiles and variation in surface molecules such as tissue factor (CD142) which may influence their function. These variations highlight the need for standardized systems to enable meaningful comparison between MSCs products.^13^

Immune checkpoint pathways, particularly PD-1 signaling, have emerged as key regulators of MSCs and T cell interactions ^14^. PD-1 engagement by its ligands PD-L1 and PD-L2 attenuates T cell activation and functions, however, the relationship between checkpoint modulation and quantitative potency readouts remains unclear ^15,16^.

In this study, we established a standardized MLR platform by optimizing key experimental variables to define a reproducible activation window. Using this system, we compared bone marrow and adipose derived MSCs across donor batches, evaluated T cell activation dynamics, and identified modulation of late activation states as a sensitive functional correlate. We further demonstrate that MSCs-mediated immunosuppression is associated with activation of a PD-1/PD-L2 regulatory axis linked to functional potency, providing a strong rationale for incorporating this mechanstic marker into standarized potency assays and future international MSCs release criteria.

## Methods

### PBMC preparation

Six buffy coats from healthy donors were obtained from Karolinska University Hospital. PBMCs were isolated using Ficoll-Paque™ PLUS (Cytiva, Sweden) and labelled with CellTrace™ Violet (CTV) Cell Proliferation Kit (Thermofisher, USA) according to the manufacturer’s instructions. Labelled PBMCs were adjusted to 2 × 10^6^ cells/mL in RPMI 1640 Gibco, USA) supplemented by 10% of USDA approval Fatal Bovin Serum (FBS) (Gibco, USA). From six individual donors, fifteen different combinations of four PBMC mixtures were prepared by combining equal volumes of cells from each donor to assess variability in T cell proliferation between different donor mixtures **(Supp table.1a)**. In parallel, a six donor PBMC mixture was prepared to evaluate whether increasing donor diversity influenced the proliferative response of T cells. To assess the impact of culture supplements on T cell proliferation, PBMC mixtures were cultured in RPMI 1640 media supplemented with (i) USDA-approved FBS, (ii) research use only (RUO) FBS, or (iii) human platelet lysate (HPL). Since MSCs were expanded in medium supplemented with 5% HPL, this condition was included for comparison. For co-culture experiments, MSCs from passage three were used. A total of 2 × 10^5^ mixed PBMCs were seeded per well in 96-well V-bottom plates and cultured for seven days. Medium exchange was performed on day three by replacing 80 µL of culture medium with 100 µL of fresh medium.

### MSCs preparation

MSCs were isolated from healthy donors following informed consent, including four bone marrow and four adipose tissue samples. Mononuclear cells (MNCs) were separated and AD derived MNCs were cultured in Minimum Essential Medium Alpha (MEM-α; Gibco, USA), while BM derived MSCs were cultured in Dulbecco’s Modified Eagle Medium (DMEM; Gibco, USA). Both media were supplemented with 5% human platelet lysate (PL Bioscience GmbH, Germany). Cells were maintained under standard culture conditions and passaged upon reaching approximately 80% confluence. MSCs were characterized according to ISCT criteria based on our previous article ^1^. All cell preparations were cryopreserved at passage three. The number of samples and production batches is summarized in **Supp table.1b**.

### Mixed Lymphocyte Reaction (MLR)

Cryopreserved MSCs from passage three were used for all experiments. AD MSCs from four donors (eight batches) and BM MSCs from four donors (nine batches) were thawed and washed in RPMI 1640 supplemented with 10% USDA approved FBS. Cells were resuspended at a concentration of 5 × 10^5^ cells/mL prior to co-culture.

MSCs were co-cultured with PBMC mixtures at MSCs:PBMC ratios of 1:4, 1:8, and 1:16 in triplicate in 96 well V bottom plates, with a final medium volume of 200 µL per well. Cultures were maintained for seven days. Unstimulated PBMCs and stimulated PBMCs without MSCs cultured as negative and positive controls, respectively. On day three, partial medium exchange was performed by removing 80 µL of culture supernatant and replacing it with 100 µL of fresh medium. At day seven the supernatant wea collected for the Elisa tests and cells were stained for analysis by FACS **(Fig.1a)**.

**Figure 1:**
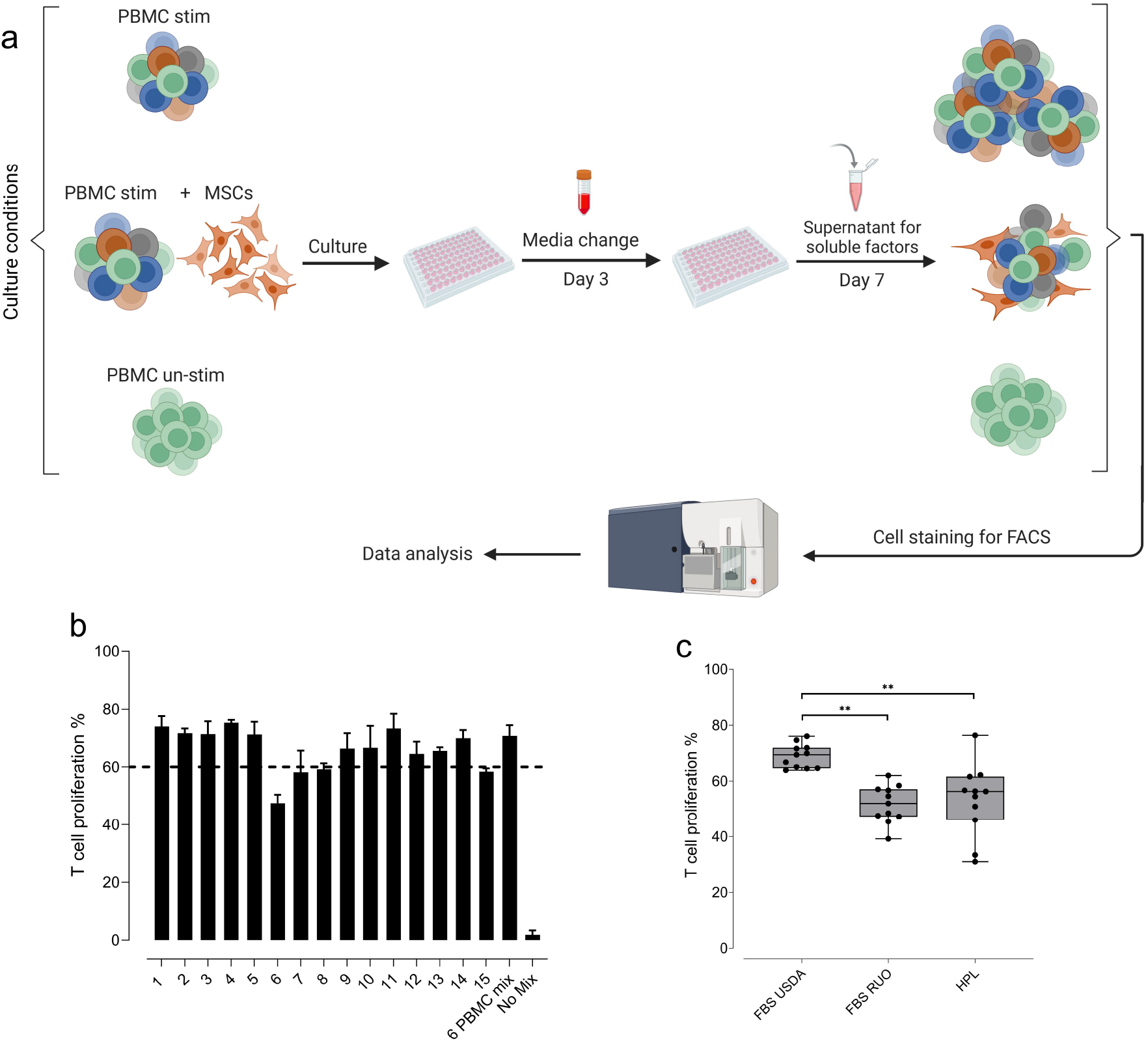
PBMC and MSCs Co-culture Workflow and Functional Assessment of T Cell Proliferation. (a) Schematic overview of the PBMC MSCs co-culture experimental workflow. (b) T cell proliferation (%) across four different PBMC mixtures and a combined mixture of six donors. The dashed horizontal line indicates the 60% threshold. (c) T cell proliferation (%) in PBMC mixtures cultured with different supplements, shown as mean and SD (^**^P ≤ 0.01).

### Flow Cytometry Analysis

After seven days of culture, cells were stained with FITC conjugated anti human CD3 (eBioscience, USA) and Fixable Viability Dye eFluor™ 780 (Invitrogen, USA). Cells were subsequently fixed using Cytofix/Cytoperm solution (BD Biosciences, USA). T-cell proliferation was assessed based on dilution of CTV within single viable CD3^+^ T cells **(Supp Fig.1)**. Samples were run in a CytoFLEX flow cytometer, and data were analysed with FlowJo software (v10.0).

To minimize inter experimental variability, T cell proliferation was normalized to the proliferation rate observed in the PBMC mixture control within each independent experiment. The suppressive effect of MSCs was calculated as the percentage of Normalized T cell Proliferation Reduction (%NTPR) using the following formula:

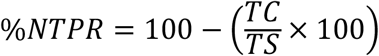

where *TC* represents the percentage of proliferating T cells in co-culture with MSCs, and *TS* represents the percentage of proliferating T cells in the stimulated PBMC mixture control.

### CD142 (TF) and Podoplanin detection

The surface expression of CD142 (tissue factor, TF) and podoplanin on MSCs was assessed by flow cytometry. Cells were stained with PE conjugated anti human CD142 antibody (eBioscience, USA) and APC conjugated anti human podoplanin (eBioscience, USA). Isotype controls and unstained cells were used to exclude non-specific binding and background fluorescence respectively.

### Detection of CD4, CD8, CD25 and HLA DR in MLR

To evaluate MSCs mediated suppression of T cell activation and proliferation, surface expression of CD25, HLA-DR together with CD4 and CD8 was analysed within single live T cells following the MLR. Cells were stained using PE conjugated anti human CD25 (Biolegend, USA), PerCP-Cy5 conjugated anti human HLA DR (Beckton Dickinson, USA), APC conjugated anti human CD4 (Biolegend, USA) and BV605 conjugated anti human CD8 (Biolegend, USA) antibodies according to standard staining protocols.

### Soluble PD-1, PD-L1, and PD-L2 Quantification

Supernatants were collected from each mixed lymphocyte reaction (MLR) condition at the day of seven and stored at −80°C until analysis. Soluble PD-1, PD-L1, and PD-L2 concentrations were quantified using commercially available ELISA kits (Abcam, Cambridge, UK) according to the manufacturer’s instructions. Optical density (OD) was measured at 450 nm using a microplate ELISA reader (PerkinElmer, USA), and concentrations were calculated from standard curves generated in parallel with each assay. To account for inter experimental variability, PD-1 and PD-L1 concentrations in MSCs and PBMC co-culture conditions were normalized to the stimulated PBMC control. Normalized concentrations were calculated using the following formula:

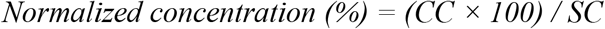

where *CC* represents the concentration of the target molecule in MSCs co cultured with stimulated PBMCs, and *SC* represents the concentration of the same molecule in the stimulated PBMC condition alone. In contrast, PD-L2 levels were reported as absolute concentrations, as PD-L2 was not detectable in the stimulated PBMC condition.

### Statistical Analysis

Data were analysed using one-way analysis of variance (ANOVA) for comparisons involving more than two group. For comparisons between two groups, an unpaired two-tailed Student’s t-test was applied. A p-value < 0.05 was considered statistically significant.

## RESULTS

### Standardization of a mixed lymphocyte reaction assay identifies key biological determinants influencing MSCs-mediated immunosuppression

To establish a robust platform for assessing the immunomodulatory potency of MSCs, we first optimized a MLR based assay modelling allogeneic immune activation following a specific protocol **(Fig. 1a)**. We first define the dynamic range of the allogeneic MLR assay by systematically evaluated T cell proliferation across multiple combinations of PBMCs derived from independent donors **(Fig. 1b and Supp table. 1a)**. Among the fifteen donor combinations tested, eleven exhibited proliferations exceeding 60%, with an average variation of less than 13% between these mixtures, indicating a reproducible activation window within this range. In contrast, four donor combinations generated lower proliferative responses ranging between 49% and 60%, suggesting a reduced allogeneic stimulation capacity. Because robust immune activation is a defining feature of highly inflammatory conditions such as GvHD or acute respiratory distress syndrome where MSCs were succesffuly used, we selected PBMC donor combinations yielding >60% proliferation as the operational threshold for subsequent assays. We also found that this cut-off threshold can be affected by the nutrient as adding another type of FBS or HPL results in a significant reduction in the proliferation of PBMCs (**Fig.1c**). Furthermore, MSCs were co-cultured with PBMCs in three different ratios. Among these, the ratio of 1:4 in MSC: PBMC demonstrated the strongest suppressive effect and was therefore selected for all subsequent experiments **(Supp Fig.1 and supp Fig. 2)**. These findings demonstrate that MLR based potency assays are highly sensitive to donor composition and culture conditions highlighting the importance of assay standardization when comparing MSCs products derived from different tissue sources or manufacturing runs.

### Quantitative potency thresholds reveal enhanced immunosuppressive activity of adipose-derived MSCs in the mix lymphocytes reactions

Using the optimized MLR system, MSCs derived from BM and AD tissues evaluated across multiple independent experimental runs. Both MSCs sources robustly suppressed T cell proliferation in stimulated PBMC cultures. However, when data from independent donor batches were analysed collectively, adipose derived MSCs consistently exhibited greater inhibitory activity than bone marrow derived MSCs under identical donor and culture conditions, indicating a reproducible difference in functional immunomodulatory potency between tissue sources **(Fig.2a)**.

**Figure 2:**
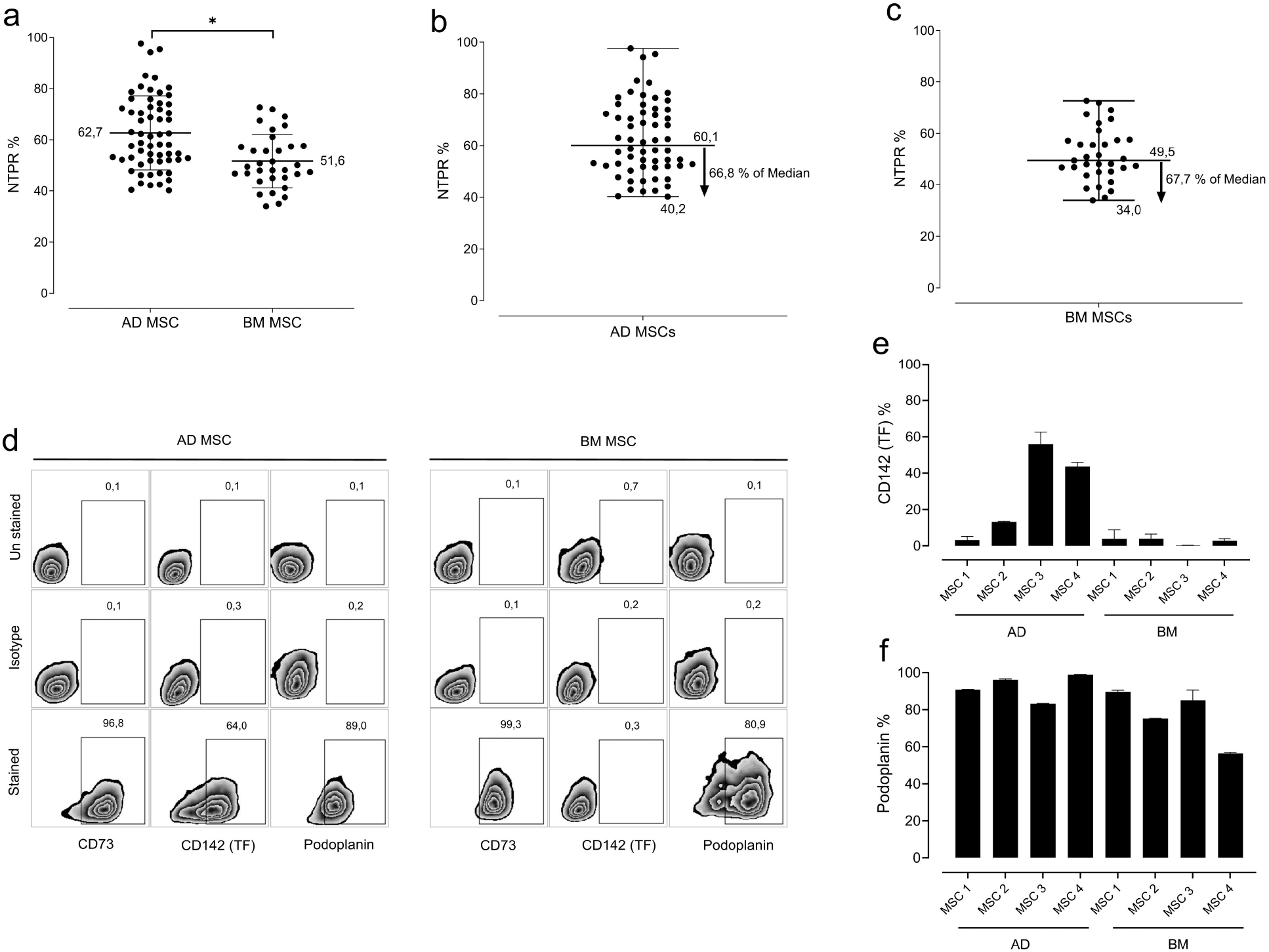
Characterization of MSCs immunomodulatory function and surface marker expression. (a) shows the percentage of NTPR in stimulated PBMCs co-cultured with AD and BM derived MSCs (mean ± SD; ^*^P ≤ 0.05). (b) and (c) graphs present the median and range of NTPR (%) for AD and BM derived MSCs, respectively. Based on the minimum suppression observed, values below the median define the lower acceptance threshold. (d) Representative flow cytometry plots showing CD142 (tissue factor, TF) and podoplanin expression on MSCs. (e) Percentage expression of CD142 (TF) and (f) podoplanin on MSCs derived from AD and BM sources.

Nevertheless, across all experimental replicates, the lowest suppression observed for adipose-derived MSCs reached 40%, whereas bone marrow–derived MSCs displayed a minimal suppression of 34%. Importantly, when normalized to the median inhibitory response within the assay, these values corresponded to a highly similar proportional reduction of approximately 68.8% and 67.7%, respectively. This convergence across independent experiments suggests the presence of a reproducible lower functional boundary of MSCs mediated immunosuppression regardless the cell origin within the MLR system. Taken together, these observations define a practical functional benchmark for MSCs potency evaluation while preserving sufficient assay sensitivity to resolve biologically meaningful differences in immunosuppressive capacity across MSC preparations **(Fig.2b and 2c)**.

Phenotypic characterization revealed that AD and BM MSCs displayed comparable expression of canonical MSCs markers included in the ISCT release criteria, including CD90, CD73, and CD105, while lacking expression of hematopoietic and endothelial markers such as CD34 and CD31. In contrast, differential expression emerged for tissue factor with CD142 affecting T cell activation and the motility marker Podoplanin **(Fig. 2d)** exhibiting distinct expression profiles between the two MSCs sources **(Fig. 2e and 2f)**. These findings suggest that while both MSCs populations conform to established identity criteria, source-dependent variation in pro-coagulant and stromal interface markers may reflect intrinsic tissue imprinting and could contribute to the observed differences in functional immunomodulatory activity.

### HLA-DR downregulation on activated T cells serves as a novel sensitive surrogate marker of MSCs functional activity

To further delineate the immunological mechanisms underlying MSCs mediated suppression within this system, we examined the expression of T cell activation markers following co-culture with MSCs. The early activation marker CD25 (IL-2Rα) and the late activation marker HLA-DR were assessed to capture distinct stages of T cell activation dynamics. To our knowledge, modulation of HLA-DR expression on stimulated T cells following MSCs exposure has not been systematically described.

In mixed lymphocyte reactions, CD25 expression was first induced following PBMC activation and was subsequently accompanied by a progressive increase in HLA-DR expression within the responding T cell population **(Fig. 3a)**. Across independent experiments, reductions in both CD25+ and CD25+ HLA-DR+ that paralleled the extent of proliferation suppression **(Fig. 3b)**. Notably, the reduction in HLA-DR expression was consistently more pronounced in cocultures containing BM MSCs compared with AD MSCs, suggesting that BM MSCs exert a stronger regulatory influence on late-stage T cell activation during allogeneic immune responses **(Fig. 3c)**.

**Figure 3:**
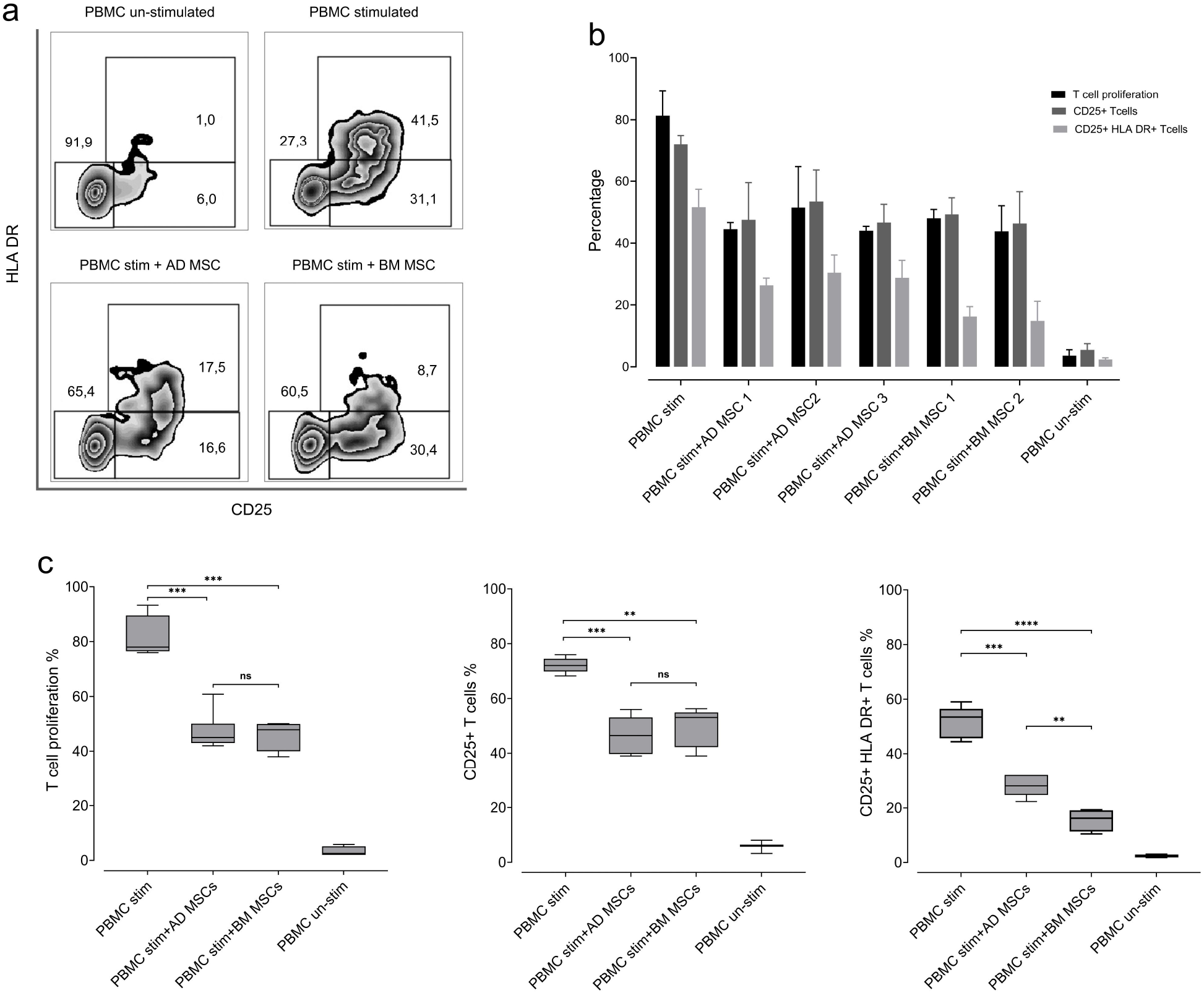
Expression of CD25 and HLA-DR in MLR conditions. (a) Representative flow cytometry density plots showing CD25 and HLA-DR expression on T cells under different conditions: stimulated PBMCs, unstimulated PBMCs, co-culture with AD MSCs, and co-culture with BM MSCs. (b) Mean percentage of proliferating T cells (based on CTV dilution), CD25^+^ T cells, and CD25^+^HLA-DR^+^ T cells in MLR. (c) Comparative analysis of T cell proliferation, CD25^+^ T cells, and CD25^+^HLA DR^+^ T cells in AD and BM derived MSCs co-cultured with stimulated PBMCs. (^**^P ≤ 0.01, ^***^P ≤ 0.001, ^****^P ≤ 0.0001)

To define the impact of MSCs mediated suppression on T cell subsets, we analysed CD4^+^ and CD8^+^ T cell populations within the MLR system. Both subsets showed robust proliferation during allogeneic stimulation **(Fig. 4a)**. Also, our data showed that in the presence of MSCs, a differential suppressive effect emerged, with a more pronounced reduction within the CD8^+^ T cell compartment. This shift was reflected in the CD4/CD8 ratio, where co-culture conditions more closely resembled the baseline ratio observed in unstimulated PBMCs **(Fig. 4b)**, suggesting preferential suppression of CD8^+^ T cell expansion.

**Figure 4:**
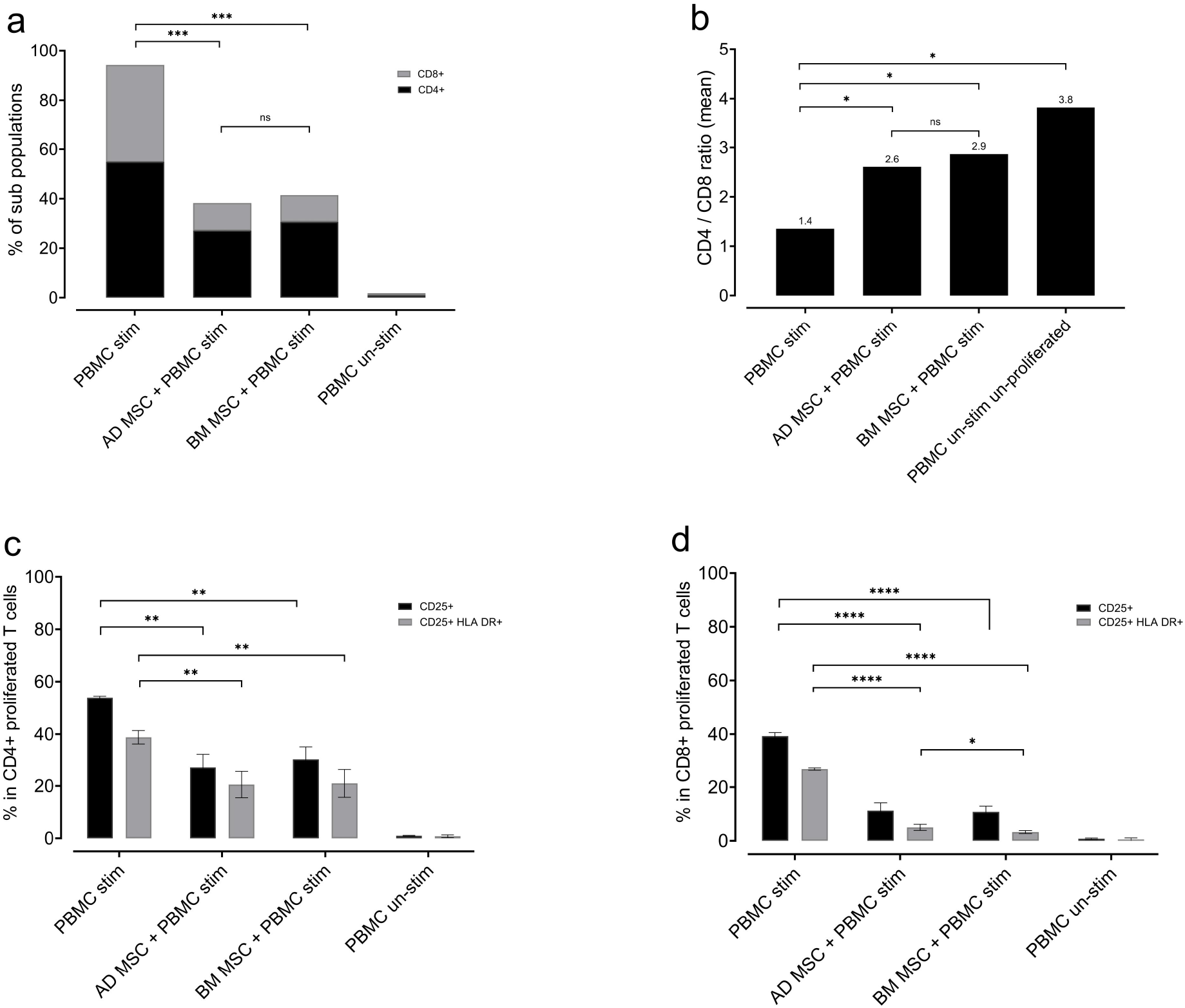
T cell subpopulations in MLR conditions. (a) Graph showing the percentages of CD4^+^ and CD8^+^ T cells under different MLR conditions. Both CD4^+^ and CD8^+^ populations are suppressed in the presence of MSCs (^***^P ≤ 0.001). (b) Ratio of CD4^+^/CD8^+^ T cells across different MLR conditions. MSCs treatment increases the CD4^+^/CD8^+^ ratio, indicating a greater suppression of activated CD8^+^ T cells (^*^P ≤ 0.05). (c, d) Expression of CD25^+^ and CD25^+^ HLA-DR^+^ T cells within CD4^+^ and CD8^+^ subsets under different MLR conditions. Both CD25^+^ and CD25^+^ HLA-DR^+^ populations are significantly reduced by MSCs, with a more pronounced effect in the CD8^+^ subset (^**^P ≤ 0.01, ^****^P ≤ 0.0001). Late-activated T cells (CD25^+^HLA-DR^+^) are further reduced by BM-MSCs compared to AD-MSCs (^*^P ≤ 0.05).

We further assessed activation markers within each subset by quantifying CD25+ and CD25^+^ HLA-DR^+^ populations. MSCs resulted in a marked reduction of both activated populations of CD4^+^ and CD8^+^ T cells, confirming effective suppression of activation in both lineages **(Fig. 4c and 4d)**. Notably, the decrease in the late activated CD25^+^ HLA-DR^+^ population was more pronounced in co-cultures containing BM MSCs compared with AD MSCs, indicating a stronger inhibitory effect of BM MSCs on late-stage T cell activation **(Fig. 4d)**.

Collectively, these findings identify downregulation of HLA-DR as a functional correlate of MSCs mediated immunosuppression and support the concept that monitoring late activation states of T cells may provide a sensitive and mechanistically informative surrogate parameter for MSCs potency assessment in MLR based assays.

### MSCs mediated immunosuppression is associated with activation of a PD-1/PD-L2 regulatory axis

To explore soluble mediators contributing to MSCs induced immunoregulation in MLR, we next examined components of the PD-1 immune checkpoint pathway. Supernatant from MSCs co-culture experiments were collected at day seven of coculture. Soluble PD-1, PD-L1, and PD-L2 were quantified in coculture supernatants by ELISA.

Activated PBMC cultures without MSCs produced detectable levels of soluble PD-1, consistent with T-cell activation. However, the addition of MSCs isolated from both bone marrow and adipose tissue resulted in a significant reduction in soluble PD-1 levels, paralleling the observed suppression of T cell proliferation **(Fig. 5a)**.

**Figure 5:**
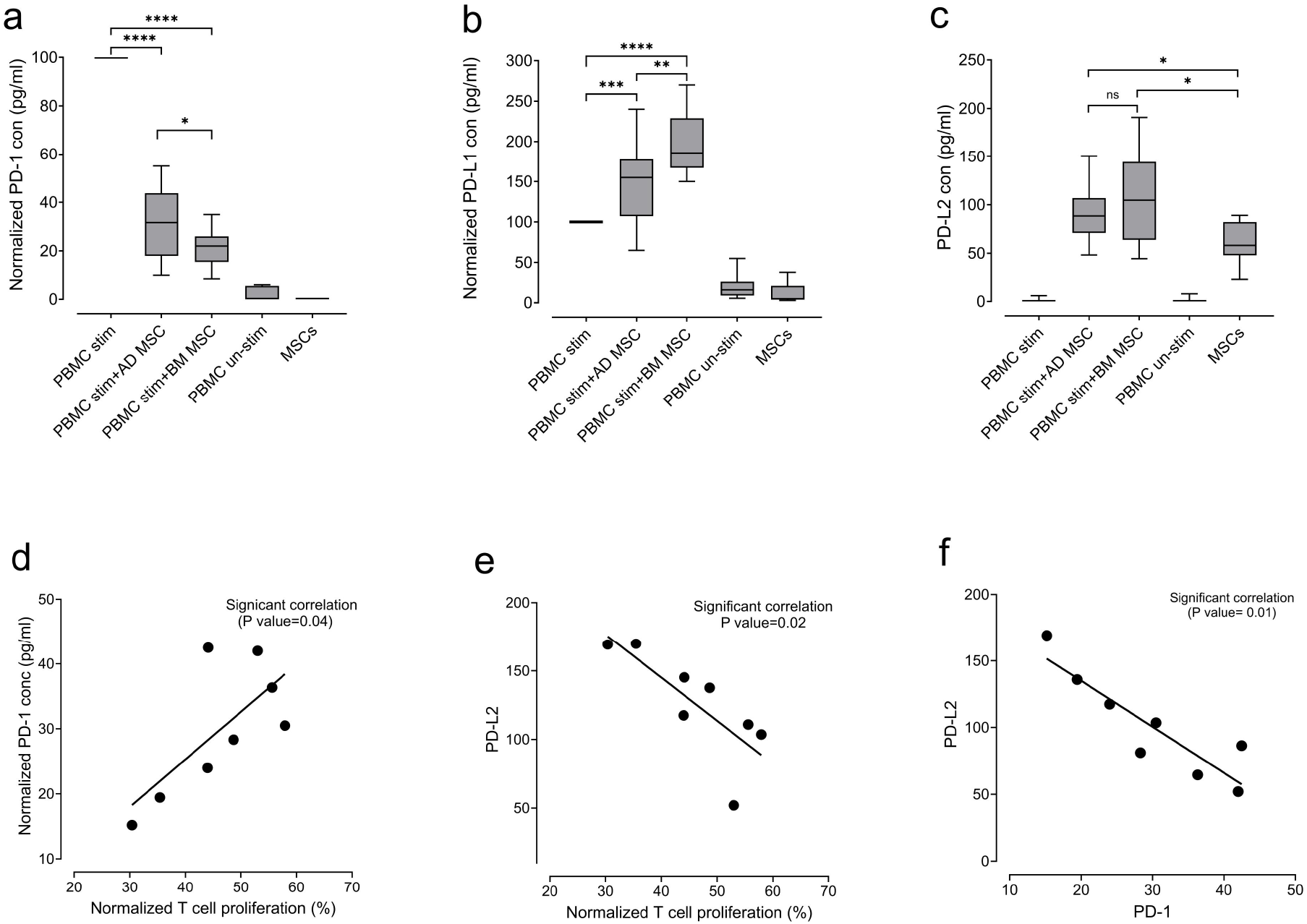
PD-1 axis levels and their correlation with T-cell proliferation in MLR. (a) concentration of soluble PD-1, (b) concentration of PD-L1, and (c) concentration of soluble PD-L2 in MLR supernatants. PD-1 and PD-L1 concentrations are normalized to the stimulated PBMC control, whereas PD-L2 is presented as an absolute concentration. (d) Correlation between PD-1 levels and T-cell proliferation, (e) correlation between PD-L2 levels and T-cell proliferation, and (f) correlation between PD-1 and PD-L2 levels. All correlations shown are statistically significant.

Moreover, soluble PD-L1 levels increased following PBMC activation and were further elevated when MSCs were present in coculture, suggesting that both immune cells and MSCs contribute to PD-L1 production in this system **(Fig. 5b)**. We observed a distinct pattern for the secretion of PD-L2. Unlike PD-L1, soluble PD-L2 was minimally detected in activated PBMC cultures alone but was readily measurable when MSCs were present. Both bone marrow and adipose derived MSCs produced PD-L2 **(Fig. 5c)** and **(Fig. 6)**. We found that the concentration of PD-L2 inversely correlated with the concentration of soluble PD-1 and of T-cell proliferation. Moreover, PD-L2 levels showed a strong inverse relationship with soluble PD-1 concentrations, suggesting coordinated regulation of the PD-1 pathway during MSCs-mediated immunosuppression **(Fig. 5d, 5e, 5f)**. However, no meaningful correlation was observed between PD-L1 levels and T-cell proliferation, nor between PD-1 and PD-L2 expression **(Supp Fig.3)**.

**Figure 6:**
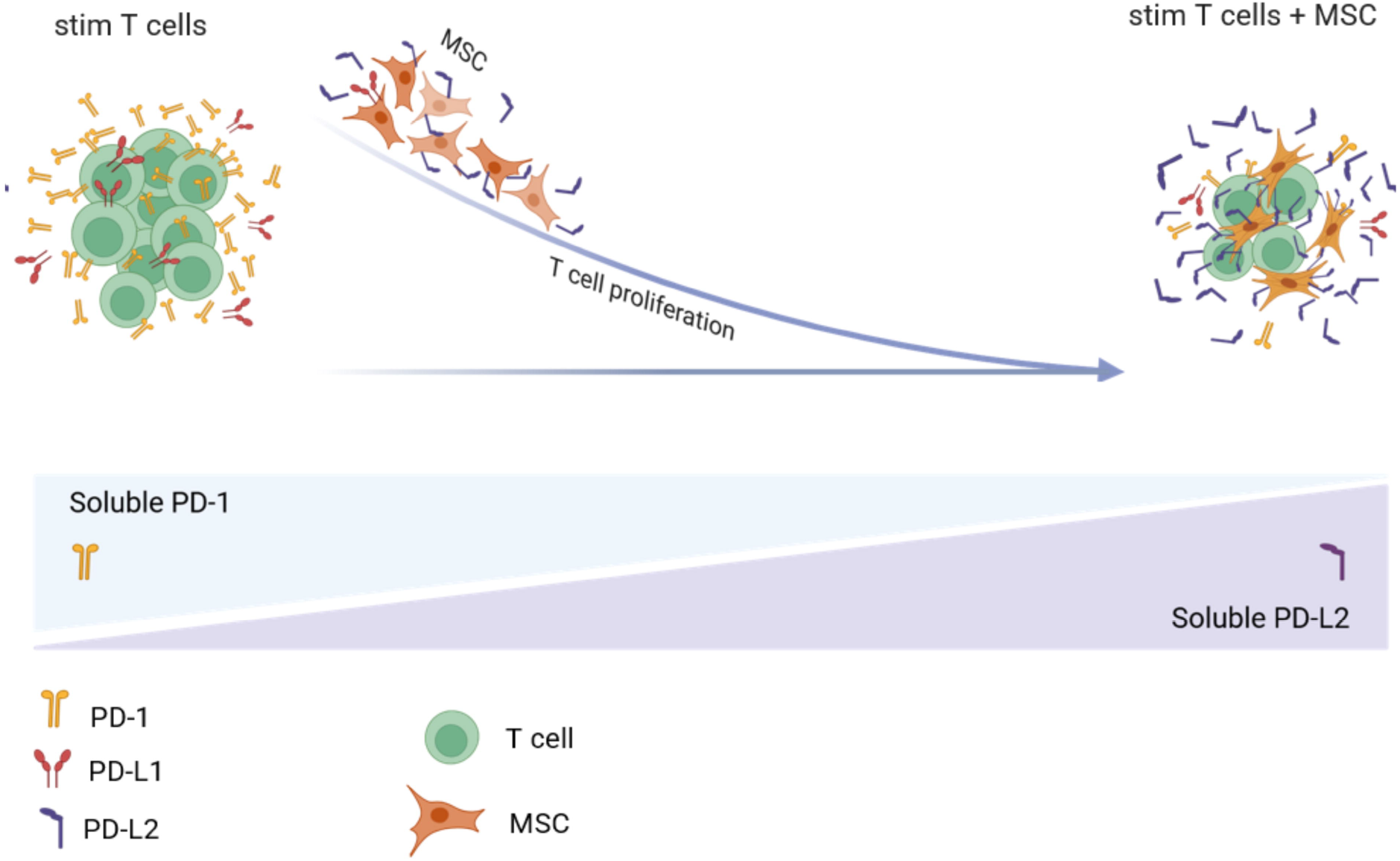
Schematic representation of PD-1 axis dynamics and their impact on T-cell proliferation in MLR.

Taken together, these findings indicate that MSCs induced immune suppression in this system is associated with modulation of the PD-1 pathway and particularly with MSC derived PD-L2 production, identifying a PD-1/PD-L2 regulatory signature that correlates with MSC functional potency.

## Discussion

In this study, we established a standardized immunosuppressive-based assay framework for assessing MSCs potency with improved robustness and mechanistic resolution. By systematically evaluating donor variability, culture conditions, and MSCs source, we demonstrate that assay configuration critically shapes functional readouts. Importantly, we define reproducible thresholds of immunosuppression, identifying modulation of HLA-DR on activated T cells as a sensitive surrogate marker, and link MSCs-mediated suppression to a PD-1/PD-L2 associated regulatory signature.

A key finding is that the dynamic range of allogeneic immune activation determines assay sensitivity. Restricting PBMC combinations to those exceeding 60% proliferation anchors the assay in a high-inflammatory state, enhancing the ability to resolve functional differences between MSCs preparations and better reflecting pathological immune contexts such as GvHD. This contrasts with conventional stimulation approaches that often obscure biologically relevant variation.^17^.

Recent efforts toward assay harmonization emphasize the need for validated, reproducible potency platforms aligned with regulatory expectations, supporting the framework proposed here ^18^. Within this system, both BM and AD MSCs exert potent immunosuppressive effects, but with distinct functional profiles. While AD MSCs demonstrate consistently higher suppression of T cell proliferation, BM MSCs preferentially attenuate late-stage T cell activation, as evidenced by more HLA-DR downregulation. This divergence suggests that tissue origin shapes distinct immunoregulatory mechanisms rather than altering the magnitude of a shared effect, consistent with known source-dependent heterogeneity. ^19^.

Despite these differences, a conserved lower boundary of immunosuppression across sources. When normalized both BM and AD MSCs exhibit a remarkably similar proportional inhibitory capacity. This observation suggesting a shared “floor effect” likely governed by common inducible suppressive mechanisms. This supports the concept that MSCs potency operates within defined biological limits rather than scaling indefinitely with dose. Consistent with this concept, prior studies have shown that MSCs-mediated inhibition of T cell proliferation reaches a plateau governed by inducible suppressive pathways such as nitric oxide or prostaglandin signalling, rather than indefinitely scaling with cell dose ^20^.

Importantly, our approach extends existing potency frameworks by defining a quantitative functional threshold within the MLR, enabling reproducible benchmarking of MSCs quality rather than relative donor ranking alone. This is in line with emerging concepts that MSCs potency is context-dependent and multidimensional. For example, Ren *et al*., showed that IL-10 induction in THP-1 cells correlates with MSCs immunosuppressive activity, where higher IL-10 responses were associated with stronger inhibition of T cell proliferation^21^. However, their approach relied on ranking donors by relative performance. Thus, rather than only identifying “more potent” versus “less potent”, our assay establishes a reproducible functional threshold allowing consistent classification of MSCs potency and improved comparability across batches. This is also conceptually consistent with Wang *et al*. who demonstrated that cytokine licensing differentially enhances MSCs immunosuppressive pathways, and with Faircloth *et al*., who showed that distinct functional outputs (angiogenesis vs immunosuppression) represent independent potency axes ^22, 23^. Together, these studies reinforce the need for context specific potency definitions, such as the immunosuppressive threshold established here.

In contrast to some earlier reports, we did not observe a correlation between PD-L1 levels and T cell proliferation. ^24^. This difference likely reflects variations in assay design. In studies such as Guan *et al*., MSCs were pre-conditioned with IFN-γ, which promotes PD-L1 expression and links it to immunosuppressive activity. In our system, immune activation occurs without artificial licensing, creating a more variable cytokine environment where PD-L1 is not the main driver of suppression. In addition, soluble PD-L1 may reflect contributions from multiple cell types rather than MSCs alone. By comparison, PD-L2 showed a stronger association with T cell suppression, suggesting it may be a more specific marker of MSCs-mediated immunoregulation in this setting.

Importantly, this distinction is further supported by differences in receptor binding properties between PD-L1 and PD-L2. PD-L2 has a higher affinity for PD-1 than PD-L1, leading to more potent inhibitory signaling ^25, 26^. This higher affinity may translate into increased functional avidity in settings of active T cell proliferation, where PD-1 expression is upregulated on activated T cells. In this context, PD-L2-mediated engagement of PD-1 may more effectively suppress proliferative responses compared to PD-L1, particularly when ligand levels are limited. Our finding that PD-L2 levels inversely correlate with T cell proliferation supports a model in which high-affinity PD-L2/PD-1 interactions preferentially regulate highly activated, PD-1 expressing T cells within the MLR system.

Taken together, these findings suggest that PD-L2 is not redundant with PD-L1 but may represent a more dominant role ligand in regulating T cell responses in physiologically relevant immune environments. This reinforces the concept that checkpoint biology in MSC-mediated immunoregulation is context-dependent and supports the inclusion of PD-L2 as a key parameter in multi dimensional potency assessment, providing a rational for its incorporation into minimal release criteria.

A key conceptual advance of this study is the identification of HLA-DR downregulation on activated T cells as a sensitive and mechanistically informative surrogate marker of MSCs activity. This dimension of T cell activation is often overlooked in MSCs studies, where CD3/CD28 driven assays dominate early activation events such as CD25, thereby underestimating the relevance of late activation markers such as HLA-DR ^27^. The consistent and pronounced reduction of HLA-DR, particularly with BM MSCs suggests that MSCs not only limit T cell expansion but also reshape activation state. Interestingly, exposure of MSCs to inflammatory environments has been shown to modulate their own HLA-DR expression, highlighting bidirectional crosstalk between MSCs and immune cells. Our findings extend this paradigm to T cell HLA-DR modulation, revealing a previously underappreciated layer of MSCs-mediated immune regulation.

At the subset level, MSCs preferentially suppressed CD8^+^ T cell proliferation, restoring the CD4/CD8 ratios toward baseline. This suggests selective control of cytotoxic T cell responses, consistent with other studies showing that pathways such as IDO can specifically inhibit CD8^+^ T cells. In this context, MSCs preferentially suppressed CD8^+^ T cells proliferation restoring CD4/CD8 balance toward baseline. This selective effect is consistent with prior work implicating metabolic pathways such as IDO in targeting cytotoxic T cell responses ^28^. Our data suggest that such mechanisms may act in concert with PD-1/PD-L2 signaling to restrain CD8^+^ T cell expansion, particularly under inflammatory conditions.

Mechanistically, our findings implicate a PD-1/PD-L2 axis as a central component of MSCs-mediated immunoregulation. Increased PD-L2 alongside reduced soluble PD-1, and its inverse correlation with T cell proliferation, support a direct role for MSCs-derived PD-L2 in suppressing T cell activation^29, 30^. Notably, our findings extend prior work by Davies et al. who demonstrated MSCs secretion of PD-1 ligands under CD3/CD28 stimulation conditions. In contrast, our MLR model captures antigen-driven activation, revealing PD-L2 as a context dependent potency marker under physiologically relevant conditions ^31^.

Taken together, these results support a multi-layered model of MSCs function integrating proliferation control, modulation of activation states, subset-specific effects, and checkpoint signaling. By defining quantifiable and mechanistically linked parameters, this work advances the development of standardized potency assays for MSC-based therapies.

Several limitations should be acknowledged. While the MLR provides a robust and controlled platform to interrogate MSCs-mediated immunomodulation, it remains an inherently reductionist *in vitro* system that cannot fully capture the complexity, spatial organization, and dynamic regulation of immune responses *in vivo*. Although functional assays are widely required by regulatory authorities for the release of MSCs based products, a consistent and predictive correlation between *in vitro* potency readouts and clinical outcomes has yet to be established. Nonetheless, such assays are essential for ensuring functional consistency under controlled conditions. Additionally, the use of soluble checkpoint measurements does not reflect cell-bound dynamics or spatial organization. Future studies integrating single-cell transcriptomics, spatial profiling, and *in vivo* validation will be essential to further refine these mechanistic insights.

In conclusion, this work provides a standardized and mechanistically anchored framework for assessing MSCs immunomodulatory potency. Identification of HLA-DR modulation, preferential CD8^+^ T cell suppression, and PD-L2 associated signalling as key functional correlates offers new tools for both research and clinical translation. These findings contribute to a more precise definition of MSC activity and support the development of next-generation potency assays capable of predicting therapeutic efficacy across diverse clinical indications.

## Acknowledgements

The authors gratefully acknowledge Cellcolabs AB for financial support of this study.

## Conflict of Interest

There is no conflict of interest.

## Author contributions

Conceptualization: NK, MNZ, Data curation: MNZ, Formal analysis: NK, MNZ Investigation: NK, MNZ, Methodology: MNZ, KL, NKH, writing original draft: NK, MNZ, Writing review and editing: NK, KL, MNZ, NKH, LS, KLB

## Figure legendss

**Supplementary Table 1. PBMC and MSCs samples used in this study**.

(a) Combinations of PBMCs from different donors used to assess variability in T-cell proliferation. (b and c) List of mesenchymal stromal cells, including their sources and batch information, used in the experiments.

**Supplementary Figure 1. Representative analysis of T-cell proliferation across different ratios and controls**. (a) The distinct peak on the right of graphs corresponds to non-proliferating T cells, while successive peaks with decreasing fluorescence intensity (leftward shift) indicate proliferating T-cell generations. T-cell proliferation was quantified based on CTV dye dilution. (b) NTPR% across different MSC sources and ratios. The graph shows the highest NTPR response observed in co-cultures of MSCs with stimulated PBMCs at 1:4, 1:8 and 1:16 ratios, consistent across all MSCs samples.

**Supplementary Figure 2. Percentage of NTPR in different ratio of co-culture**. NTPR% in stimulated PBMCs cocultured with AD and BM derived MSCs (mean ± SD) in 1:4, 1:8 and 1:16 ratios.

**Supplementary Figure 3. Soluble PD-L1 level and its correlation with T-cell proliferation and PD-1 in MLR**. The graphs show that soluble PD-L1 levels do not significantly correlate with T-cell proliferation (a) or PD-1 expression (b).

## Notes

### Competing Interest Statement

The authors have declared no competing interest.

