## Supplementary figures and images for "HLA-DR MODULATION AND PD-1/PD-L2 CHECKPOINT SIGNALLING DEFINE A MECHANISTIC POTENCY AXIS FOR MESENCHYMAL STROMAL CELL IMMUNOSUPPRESSION"

### Supplementary Figure 1

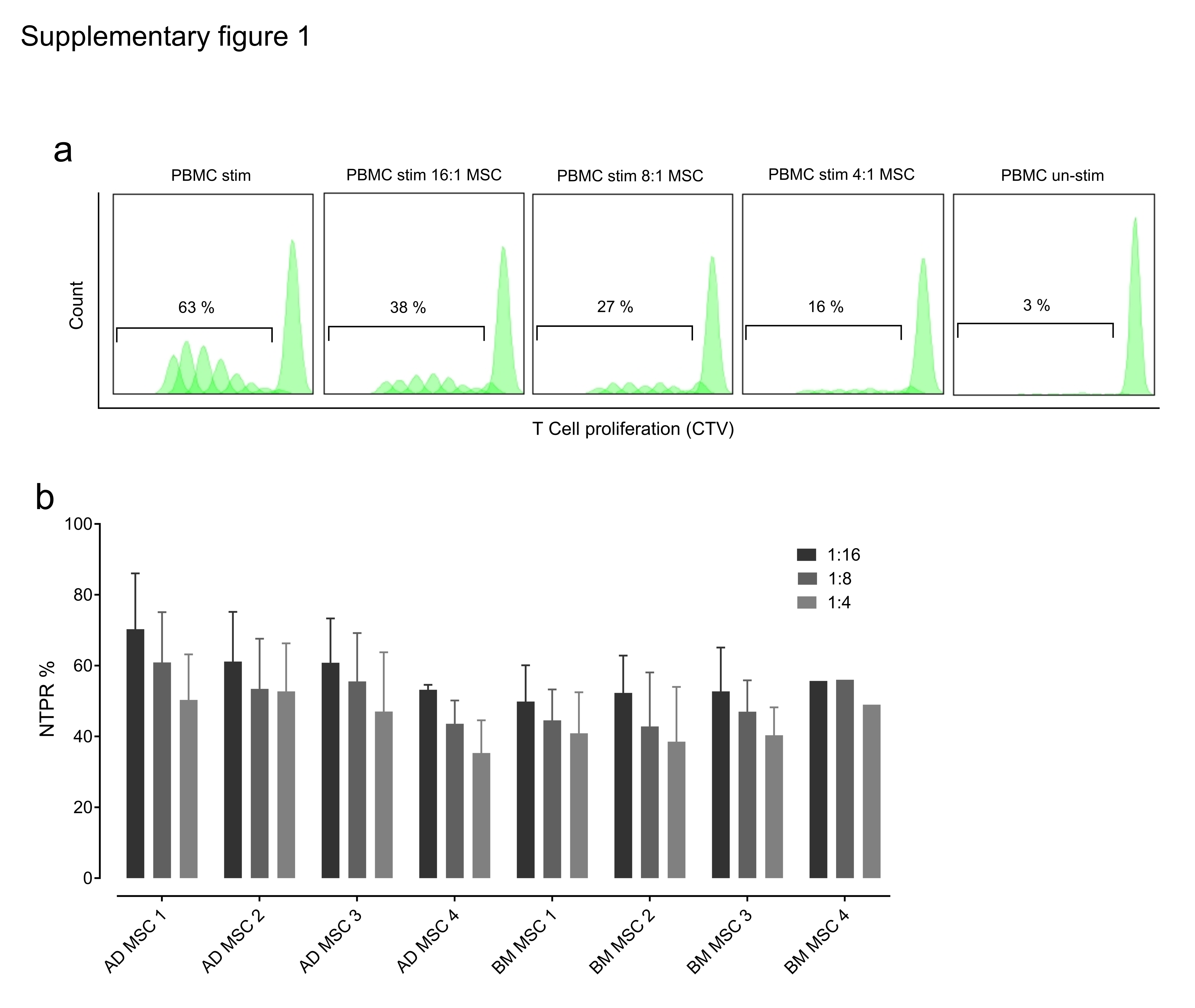

### Supplementary Figure 2,3

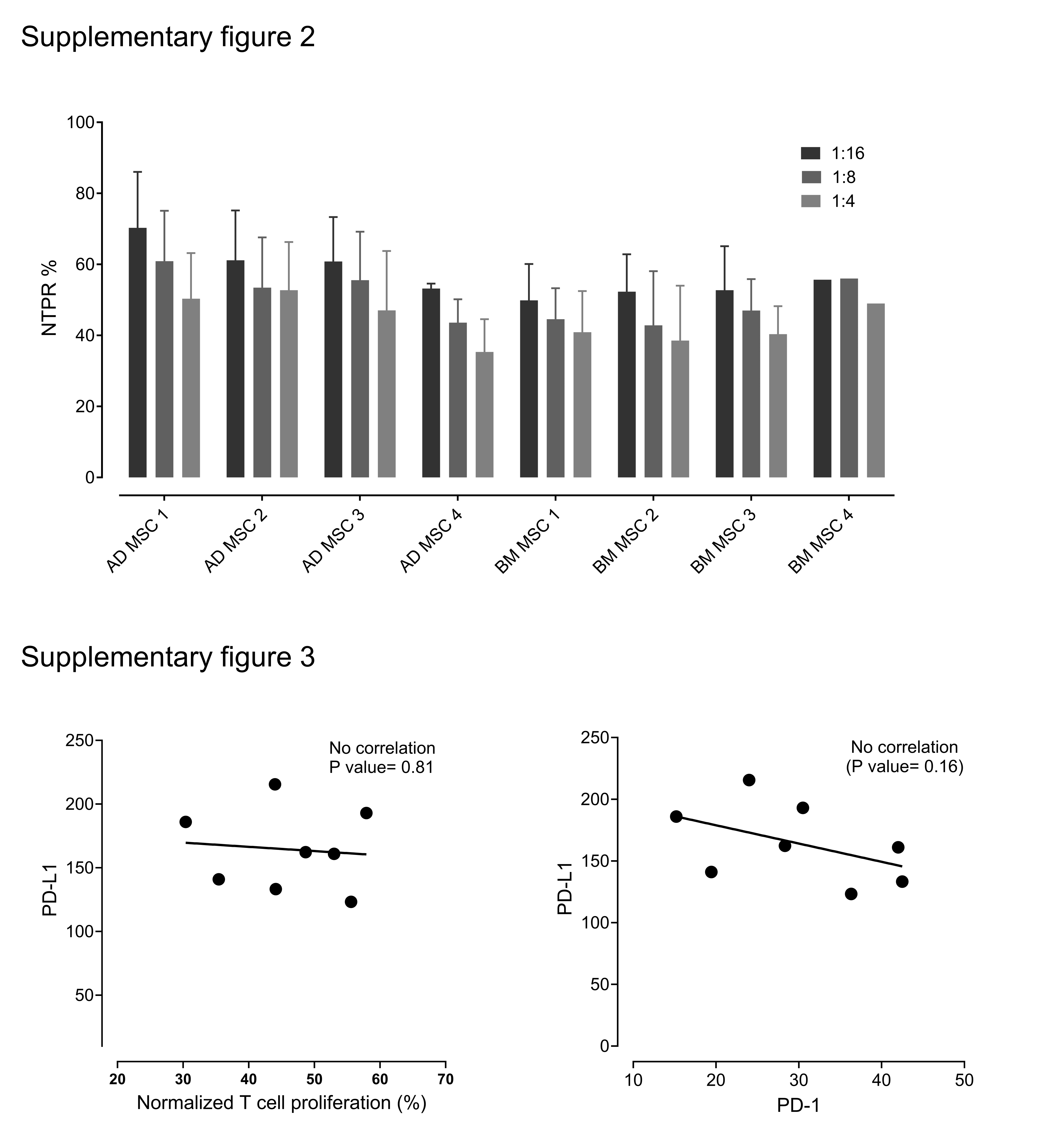

### Supplementary table 1

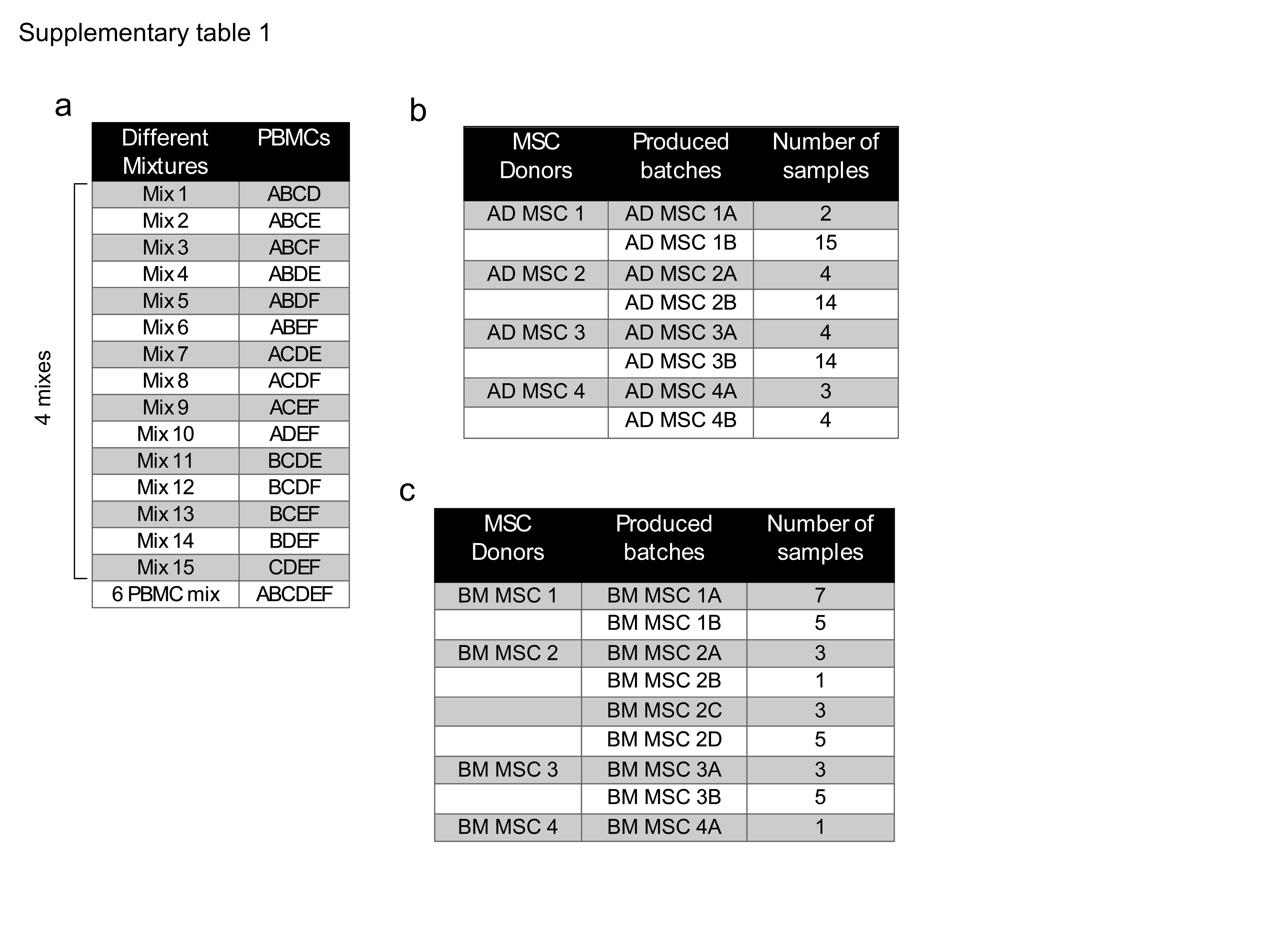
